# The Nested Hierarchy of Overt, Mouthed, and Imagined Speech Activity Evident in Intracranial Recordings

**DOI:** 10.1101/2022.08.04.502829

**Authors:** Pedram Z. Soroush, Christian Herff, Stephanie K. Ries, Jerry J. Shih, Tanja Schultz, Dean J. Krusienski

## Abstract

Recent studies have demonstrated that it is possible to decode and synthesize various aspects of acoustic speech directly from intracranial measurements of electrophysiological brain activity. In order to continue progressing toward the development of a practical speech neuroprosthesis for the individuals with speech impairments, better understanding and modeling of imagined speech processes are required. The present study uses intracranial brain recordings from participants that performed a speaking task with trials consisting of overt, mouthed, and imagined speech, representing various degrees of decreasing behavioral output. Speech activity detection models are constructed using spatial, spectral, and temporal brain activity features, and the features and model performances are characterized and compared across the three degrees of behavioral output. The results indicate there is a hierarchy in which the relevant channels for the lower behavioral output modes form nested subsets of the relevant channels from the higher behavioral output modes. This provides important insights for the elusive goal of developing more effective imagined speech decoding models with respect to the better-established overt speech decoding counterparts.

## I. Introduction

Speech is the first and foremost means of human communication. Millions of people worldwide suffer from severe speech disorders due to neurological diseases such as amyotrophic lateral sclerosis (ALS), brain stem stroke, and severe paralysis. A speech neuroprosthesis that decodes speech directly from neural signals could dramatically improve life for these individuals. Intracranial brain-computer interfaces (BCIs) using electrocorticography (ECoG) [1], [2], [3], [4], [5], [6], [7], [8] or stereotactic electroencephalography (sEEG) [2], [8], [9], [10], [11], [12] have demonstrated success in decoding aspects of speech directly from brain activity.

For those who have completely lost the ability to speak, the objective is to synthesize acoustic speech directly from brain activity during *imagined* speech. However, the lack of acoustic or behavioral output during imagined speech presents challenges in designing an effective decoding model [8], [13], [14], [15]. To cope with this limitation, it is common to utilize behavioral output from *overt* speech or *mouthed* speech (i.e., performing inaudible speaking articulations without vocalization) as a surrogate to study associated brain activity [3], [4], [5], [16], [17], [18], [19], [20] or to train decoding models for imagined speech applications [8], [21], [22].

Numerous prior studies have focused on establishing neural speech decoding model performance in various scenarios such as decoding phonemes and words [16], [23], [24], brain-to-text [3], [6], [7], [25], [26], and direct speech synthesis from brain activity [4], [5], [8], [22]. However, few studies have systematically compared the characteristics of the decoding models across overt, mouthed, and imagined speech modes. In order to simplify the problem for such a comparative study, a speech activity detection framework can be utilized [27], [28], [29]. Using causal brain activity features as inputs, speech activity detection models simply classify the respective activity as occurring during intervals of *speech* or *non-speech* intent, whether overt or imagined. Recent studies using sEEG and speech activity detection modeling have successfully elucidated relevant frequency bands’ relative contributions of grey and white matter over speech [2] and classified phonetic features from activity located in the superior temporal gyrus [30].

The present study utilizes sEEG recordings, spanning cortical and subcortical areas, to compare brain activity during overt, mouthed, and imagined speech with the objective of elucidating the efficacy of various sEEG features and speech surrogates for informing imagined-speech decoding models. Multiple speech activity detection decoding models are developed and applied within and across overt, mouthed, and imagined speech modes to reveal the similarities and differences in the relevant spatial, temporal, and spectral features among the models. The results indicate that relevant channels for speech decoding reside in both cortical and subcortical areas and appear to form a hierarchy in which the relevant channels for the lower behavioral output modes form nested subsets of the relevant channels from the higher behavioral output modes.

## II. Methodology

### A. Participants and Electrode Locations

sEEG data were collected from 7 native English-speaking participants being monitored as part of treatment for intractable epilepsy at UCSD Health. The demographic information of the participants is provided in Table I. The study design was approved by the Institutional Review Boards of Virginia Commonwealth University and UCSD Health, and informed consent was obtained for experimentation with human subjects. The locations of sEEG electrodes were determined solely based on the participants’ clinical needs. A subset of the implanted electrodes for each participant was determined to be in or adjacent to brain regions associated with speech and language processing. Fig. 1 shows the depth electrode locations for the 7 participants, with sEEG electrode (channel) counts provided in Table I. Anatomical location of the channels, including brain region and localization in white or grey matter, were identified using the FreeSurfer software package and MNE-Python [31], [32].

**TABLE I:**
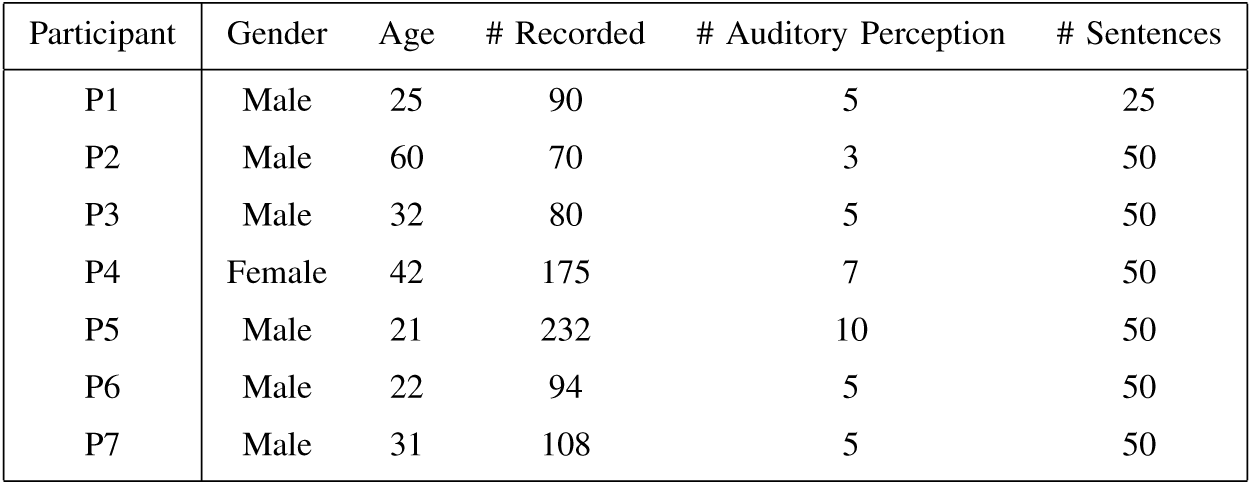
Demographic information of participants and numbers of sEEG channels and unique Harvard sentence task sequences performed for each participant. The first and second columns list the gender and age of the participants, respectively. The third column reports the total number of channels recorded during the experiment, and the fourth column reports the subset of channels identified as located in auditory regions and having potential auditory perception information as described in Section II-F. The last column lists the number of Harvard sentence task sequences performed.

| Participant | Gender | Age | # Recorded | # Auditory Perception | # Sentences |
| --- | --- | --- | --- | --- | --- |
| P1 | Male | 25 | 90 | 5 | 25 |
| P2 | Male | 60 | 70 | 3 | 50 |
| P3 | Male | 32 | 80 | 5 | 50 |
| P4 | Female | 42 | 175 | 7 | 50 |
| P5 | Male | 21 | 232 | 10 | 50 |
| P6 | Male | 22 | 94 | 5 | 50 |
| P7 | Male | 31 | 108 | 5 | 50 |

**Fig. 1:**
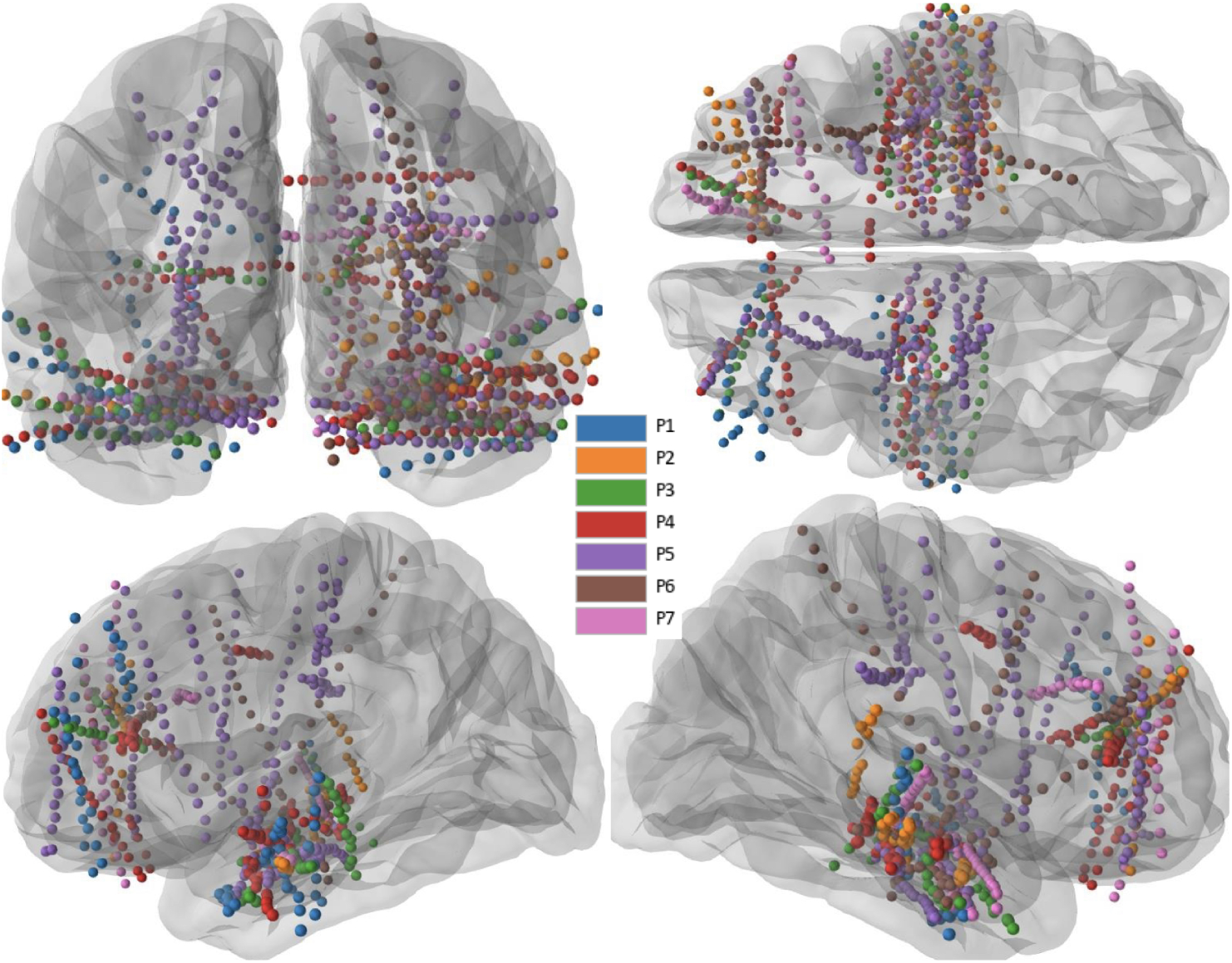
The combined sEEG depth electrode (channel) locations of the 7 participants from different perspectives using an averaged brain model.

### B. Experimental Design

For the experiment, participants were presented with a sentence displayed on a computer monitor and simultaneously narrated via computer speakers. When visually prompted with an icon as shown in Fig. 2a, the participant was instructed to speak the sentence audibly while the acoustic speech and sEEG signals were simultaneously recorded. The participant was subsequently visually prompted with icons indicating to inaudibly articulate speech (i.e., mouth) and imagine speaking without articulating or vocalizing, respectively, for the same sentences. Herein, these three modes are referred to as *overt, mouthed*, and *imagined*, respectively. Each icon prompt and participant response during the task is referred to as a single *trial*.

**Fig. 2:**
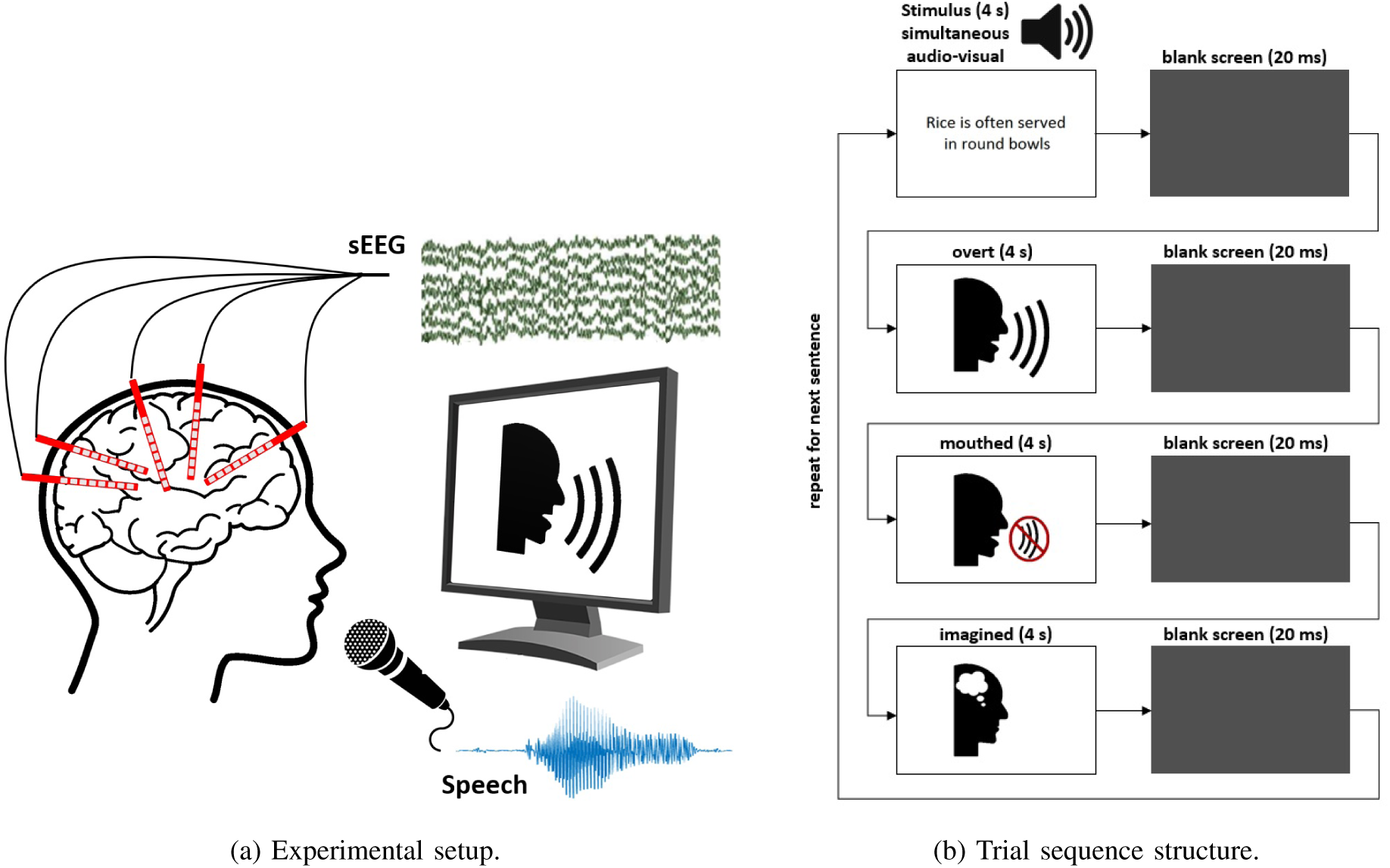
(a) sEEG and acoustic speech were simultaneously recorded as the participant performed the task as prompted by icons presented on a monitor. (b) Participants were presented with a sentence displayed on a monitor and simultaneously narrated through speakers. Participants were instructed to speak (overt), mouth, and imagine the sentence in sequence as cued by the respective icons. This was repeated for a bank of 50 sentences.

The participant was asked to perform the associated task immediately upon presentation of the icon within a 4-second interval. This structure was repeated for 25-50 unique Harvard sentences, which are phonetically-balanced based on conversational English [33].

The three tasks were intentionally presented in a consistent sequence according to degree of behavioral output (i.e., overt-mouthed-imagined) rather than block randomized to better facilitate compliance from the patients. It is believed that the behavioral output of the speaking trial better primes the participant for more reliably performing the subsequent mouthed and imagined trials. Moreover, randomizing the trials would require greater attentional resources and more likely lead to response errors and oddball effects [34], particularly with this patient group who are sometimes attentionally compromised due to their condition and the stress of the hospital environment.

The number of unique sentences for each participant is provided in Table I. The stimuli were presented and synchronized with the sEEG recordings using Presentation® software (Version 18.0, Neurobehavioral Systems, Inc., Berkeley, CA, www.neurobs.com). The experimental setup and trial sequence structure are depicted in Fig. 2.

### C. Data Collection

The sEEG electrodes (Ad-Tech Medical Instrument Corporation) were referenced to a pair of subdermal needle electrodes in the scalp and digitized at 1,024 Hz (Natus Quantum Amplifier, Natus Medical Inc.). The audio signal, recorded via an external microphone, was digitized at 44,100 Hz. The data from the audible speech portions of the task were used to extract speech and non-speech segments from the audio recordings.

### D. Labeling the Audio Files (Speech vs. Non-speech)

The recorded speech from the overt mode was manually transcribed using the Wavesurfer software package [35] for a separate analysis, but was found useful to provide precise labeling of the *speech* and *non-speech* segments for the present study. This was accomplished by shifting a 10 ms non-overlapping frame across the audio recording to identify the onset and offset of the spoken sentence, with the resulting timings from the transcription word boundaries being used as the frame label. Each frame was identified as *speech* if at least half of the frame length overlapped with a transcribed word, and as *non-speech* otherwise. For each 4-second interval encompassing the entire sentence utterance, the entire duration between the first onset and last offset was labeled as *speech* and the periods before and after these were labeled as *non-speech*. The frame length was chosen to be 10 ms to better represent brain signals’ non-stationary nature and the fast changes of speech activity for eventual closed-loop implementation.

### E. Labeling the Audio-less Modes (mouthed and imagined)

Due to non-existent speech audio for the mouthed and imagined modes, the average onset timings and durations of the overt speech intervals and the audio data from the corresponding overt mode were used to define respective surrogate ‘speech’ and ‘non-speech’ labels for the mouthed and imagined speech modes [36]. For each sentence and mode, the onset and offset of speech activity were estimated based on the average onset and offset timings of the overt mode of the corresponding participant, while the duration of speech interval was determined according to the respective overt audio. The start of each mode trial (i.e., time *t*_0_) to time *t*_1_ was labeled as *non-speech*, from time *t*_1_ to *t*_2_ was labeled as *speech*, and time *t*_2_ to the end of the trial was labeled as *non-speech*.

For each participant and sentence, the average latency between presentation of the vocalization icon and the onset of actual vocalization was computed (*t*_*a*_). This average latency was used as the transition from *non-speech* to *speech* (*t*_1_ = *t*_0_ + *t*_*a*_). To set the transition from *speech* to *non-speech* (*t*_2_), the duration of the corresponding sentence vocalization (*t*_*s*_) was used (*t*_2_ = *t*_1_ + *t*_*s*_). The interval from *t*_2_ to the end of the trial was labeled as *non-speech*.

### F. Data Pre-processing

All sEEG data were visually inspected and noisy or anomalous channels and segments were excluded from the models. Additionally, the sEEG data were analyzed for potential spatio-temporal correlations with the sound produced by the participants or present in the environment to verify the recordings were not subject to audio contamination [37]. Additionally, channels in the vicinity of the auditory cortex and belts exhibiting large relative correlations between broadband gamma activity and audio recordings of the narrated sentences were identified. These auditory perception channels could potentially contribute perceptual information to the models and misleadingly affect their performance.

To identify the auditory perception channels, the sEEG signals recorded during the audible presentation of the sentences were compared to the corresponding auditory stimulus recordings. The broadband gamma was computed for each sEEG channel using a sixth-order Butterworth IIR zero-phase filter from 70-170 Hz, notch filtered on 120 Hz. The Hilbert transform was used to compute the amplitude envelope of the resulting broadband gamma and the audio recordings. The resulting amplitude envelopes were smoothed using an eight-order Chebyshev Type I lowpass filter with cutoff frequency of 8 Hz, and decimated to 100 Hz [15]. Next, for each participant and for each audio file, the unbiased cross-correlation between the envelopes of the audio file and the envelopes of each sEEG channel were calculated for positive temporal lags up to 200 ms, and the peak of the cross-correlation was calculated and averaged through audio files for each sEEG channel. Channels exhibiting correlations above two standard deviations from the mean correlations of all channels were flagged as directly responsive to the auditory stimulus. The number of identified auditory perception channels for each participant is provided in Table I.

Because the identified auditory perception channels were found to not contribute to the resulting mouthed and imagined models in a prior analysis (following the same modeling process discussed in Section II-H), these channels were excluded from further analysis so as to not favorably and misleadingly bias the overt model results due to the availability of perceptual feedback solely for that mode. The resulting raw sEEG channels for inclusion in the analysis were re-referenced using the Laplacian method [38], [39].

### G. Feature Extraction

The narrow-band power of each sEEG channel was computed in four conventional frequency bands: theta (4-8 Hz), alpha (8-12 Hz), beta (12-30 Hz), and broadband gamma (70-170 Hz). These frequency bands have been shown to be informative for speech activity detection [2], [27], [40]. The subsequent pre-processing was performed for each frequency band.

To extract the features, using the labeled 10 ms frames from the audio signals, the sEEG channels over a specified temporal window around each audio frame were zero-phase filtered over the respective frequency range using a sixth-order Butterworth filter. The window length was chosen to be 210 ms (corresponding to 200 ms before the frame to the end of the frame). However, this is insufficient for estimating the lower frequency bands as at least 3-4 cycles are needed to convey meaningful information in a particular band. Hence, for theta, alpha, and beta bands, the duration of four cycles of the lowest frequency of the band was used to determine the causal model’s window onset, and the window offset was always fixed at the 10 ms frame length. For example, for the 4-8 Hz theta band, the window onset is 1 second (4 cycles *×* 0.25 s/cycle) before the start of the frame, giving a 1.01 s window length.

For each channel and frequency band, each temporal window was band-pass filtered over the frequency range of the respective band using a sixth-order Butterworth zero-phase IIR filter. An additional 118-122 Hz notch filter was applied to broadband gamma to suppress the second harmonic of the 60 Hz line noise. Finally, the features were computed every 10 ms as the natural logarithm of the signal energy over 210 ms, representing 10 ms overlapping the audio frame and 200 ms prior to the frame to emulate a causal design. Such a causal design aims to decode activity related to speech production rather than perception and can be implemented for real-time feedback for future closed-loop applications. The features from each included channel were concatenated to form the feature vector (# channels *×* 21 features *×* # frequency bands) for the decoding models. A diagram of the feature extraction process is provided in Fig. 3.

**Fig. 3:**
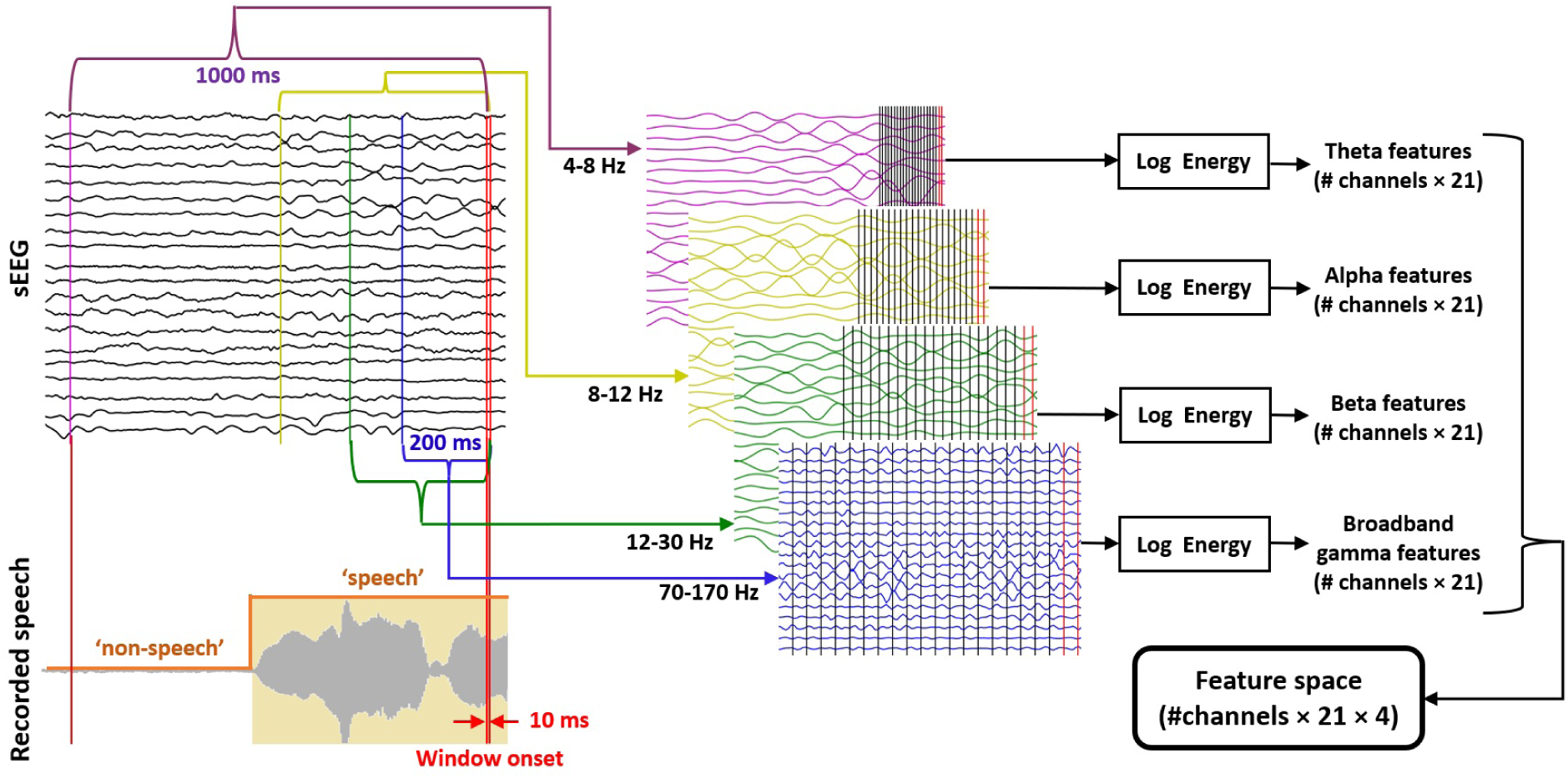
Extraction of theta, alpha, beta, and broadband gamma features from 200 ms prior to the audio frame to the frame’s offset, resulting in a feature space of # channels *×* 21 temporal lags *×* 4 frequency bands. The recorded speech is time-aligned with the sEEG and indicator of the ‘speech’/’non-speech’ labeling is shown. The filtered bands are presented in different time scales, with the vertical bars indicating the 21 10-ms temporal frames for each band.

### H. Model Training and Evaluation

All significance tests were performed using a Benjamini-Hochberg corrected Wilcoxon signed-rank test [41]. The resulting p-value (*p*) and sample sizes (*n*) are reported for each respective test.

#### 1) Logistic Regression Model

All models are designed using logistic regression with L1 regularization and are specific for each participant and mode. A proximal ADAGRAD optimizer with SoftMax function was selected for training the model [42]. This model was chosen over more complex machine learning and deep learning models because of the small size of data available (due to the differences between brain coverage of different participants, data from individual participants is processed separately), and that it has been shown to be effective for evaluating individual features and providing a convenient interpretation of the feature contributions [2].

A 10-fold cross-validation analysis was employed. To prevent training bias, the training data were normalized to zero mean and unit variance, and the same normalization parameters were applied to the test data. For each model, one tenth of the training data was used as a validation set to optimize the hyperparameters of the training models. Due to the difference between the amount of data for each class in some of the models, and to have consistency, the performance of all models was evaluated using balanced accuracy.

For all models, the balanced accuracy was evaluated as the average of the recalls of the classes, which ranges from 0 (i.e., worst possible performance) to 1 (i.e., best possible performance). To establish the chance-level classification, a randomization test was performed where all labels were randomly shuffled and the 10-fold cross-validation process was repeated for 1,000 separate randomizations of the labels.

#### 2) Single-Channel Models and Channel Selection

Single-channel decoding models were created to compare the relative decoding performance of each sEEG channel for a given mode, referred to as Within-Mode (WM) models. For each mode, the decoding performance was evaluated using 10-fold non-shuffled cross-validation models. Additionally, the feature weights of the 10 decoding models (one per fold) were averaged to compare the relative contribution of each feature to the models. These weights can provide a convenient interpretation of individual feature contributions based on the non-zero classifier weights and represent the spectral and temporal contributions to speech activity detection [2].

The single-channel WM models were subsequently used to select channels with relatively superior performance in comparison with the rest of the channels. For each mode and fold in the cross-validation process, the mean plus one standard deviation of the balanced accuracy of all channels was determined as the threshold for the fold to form a distribution of thresholds over the ten fold of the cross-validation process. Additionally, for each channel, the distribution of balanced accuracies over the 10 folds of the cross-validation process was computed.

Single-channel Cross-Mode (CM) models were also trained on the data from one mode (train mode) and respectively tested on the other two modes (test mode). The hyperparameters of the decoding model were selected based on the single-channel WM model of the train-mode. The purpose of these models is to identify channels that exhibit similar relevant features across various modes versus channels that have dominant features that are unique to specific modes.

In a separate analysis using the data from the forty channels that were excluded in the auditory perception pre-screening processes, single-channel WM models were trained and their performances were compared with the rest of the non-auditory channels. Out of the forty channels, more than thirty five channels had significantly above threshold performance for overt, but only seven and three exhibited significantly above threshold performance for mouthed and imagined, respectively. Thus, all the channels identified in the auditory perception pre-screening were excluded from the analysis.

#### 3) Multi-channel Models

Using the features of all channels, for each participant and mode, multi-channel WM models were evaluated using 10-fold non-shuffled cross-validation process. Multi-channel CM models were also created for which a decoding model was trained on the entire data of the train-mode and tested on the entire data from a different test-mode, herein labeled as (train mode)-to-(test mode). The hyperparameters of the decoding model were selected based on the multi-channel WM model of the train-mode. The purpose of these models is to identify similarities and differences in the decoding models and performance across modes.

## III. Results

The three modes can be compared with respect to degree of behavioral output, with overt having the highest, mouthed having intermediate, and imagined having no behavioral output. In the subsequent paired comparisons, the mode in the pair having the higher behavioral output will be denoted as the *HBO mode* and the mode with lower behavioral output will be denoted as the *LBO mode*.

### A. Models

Fig. 4 illustrates violin plots of the distributions of averaged balanced accuracy of the 10-fold cross-validation single-channel WM models trained on the data from each participant and each mode. The black dots represent the channels that performed significantly above the thresholds (*p <* 0.01, *n* = 10) and were thus selected as mode-relevant channels. For all participants and modes, the thresholds were significantly above the chance-level, which was approximately 0.5 on average, (*p <* 0.01, *n* = 10). Additionally, over all participants, channels that performed significantly better than chance-level (*p <* 0.05, *n* = 10) in all three modes performed significantly better in overt compared to mouthed and imagined modes, and in mouthed compared to imagined (*p <* 0.01, *n* = 10).

**Fig. 4:**
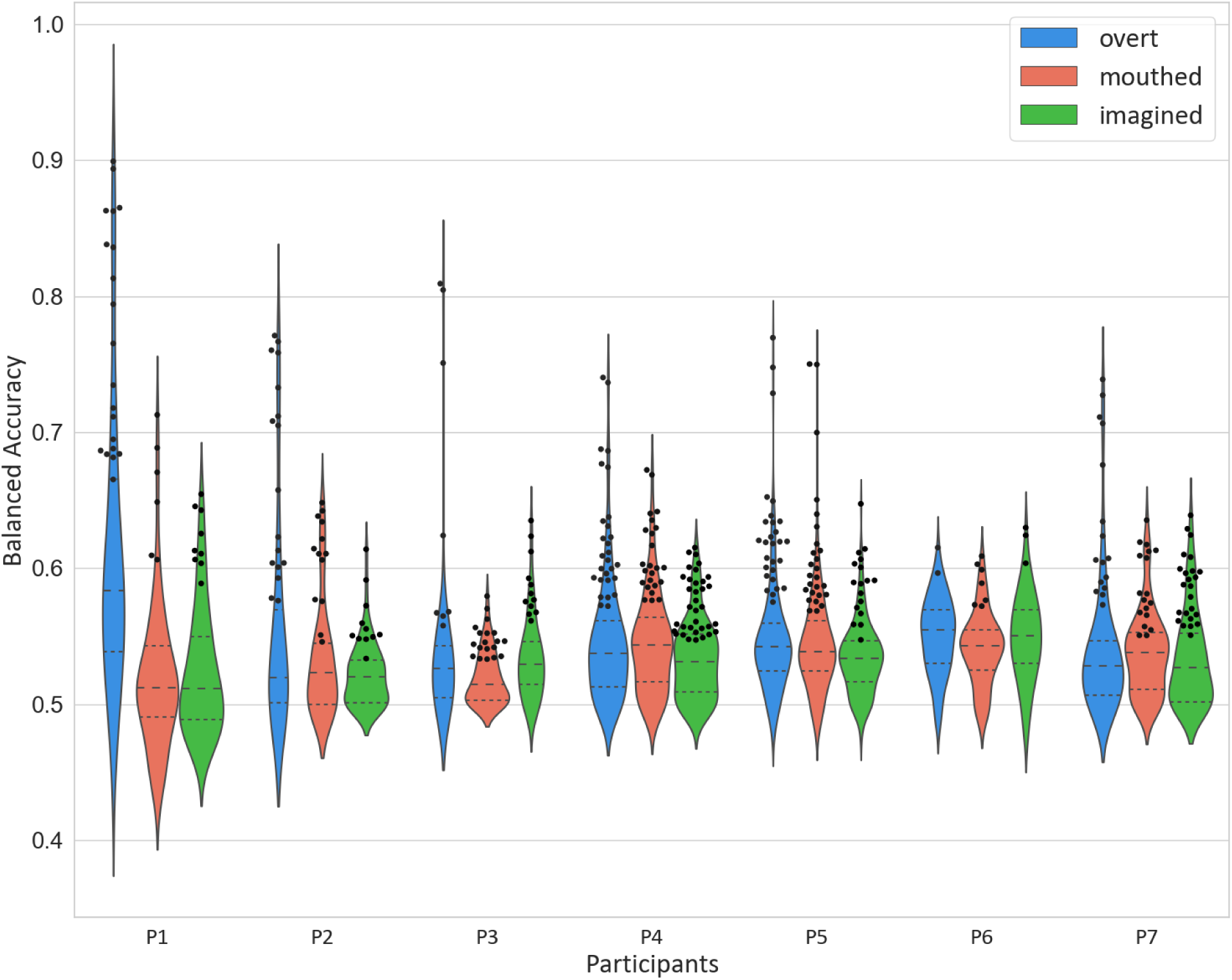
Distributions of classification performance (averaged balanced accuracy over 10-fold non-shuffled cross-validation models) of decoding models for all channels in the single-channel WM models for each participant and mode. The black dots represent the selected channels for each group. The dashed line within each violin indicate the median and the dotted lines indicate the first (Q1) and third (Q3) quartiles. Chance-level was approximately 0.5 on average.

#### 1) Spatial Characterization

Fig. 5 shows the individualized averaged balanced accuracy of the cross-validated single-channel WM models on the data from a representative participant (Participant 1) for all three modes. This is equivalent to an expansion of the violin plots from Participant 1 in Fig. 4. Fig. 5 shows that channels along each shaft, if relevant for the mode and significantly above the chance-level (*p <* 0.05, *n* = 10), generally follow a similar trend across the modes.

**Fig. 5:**
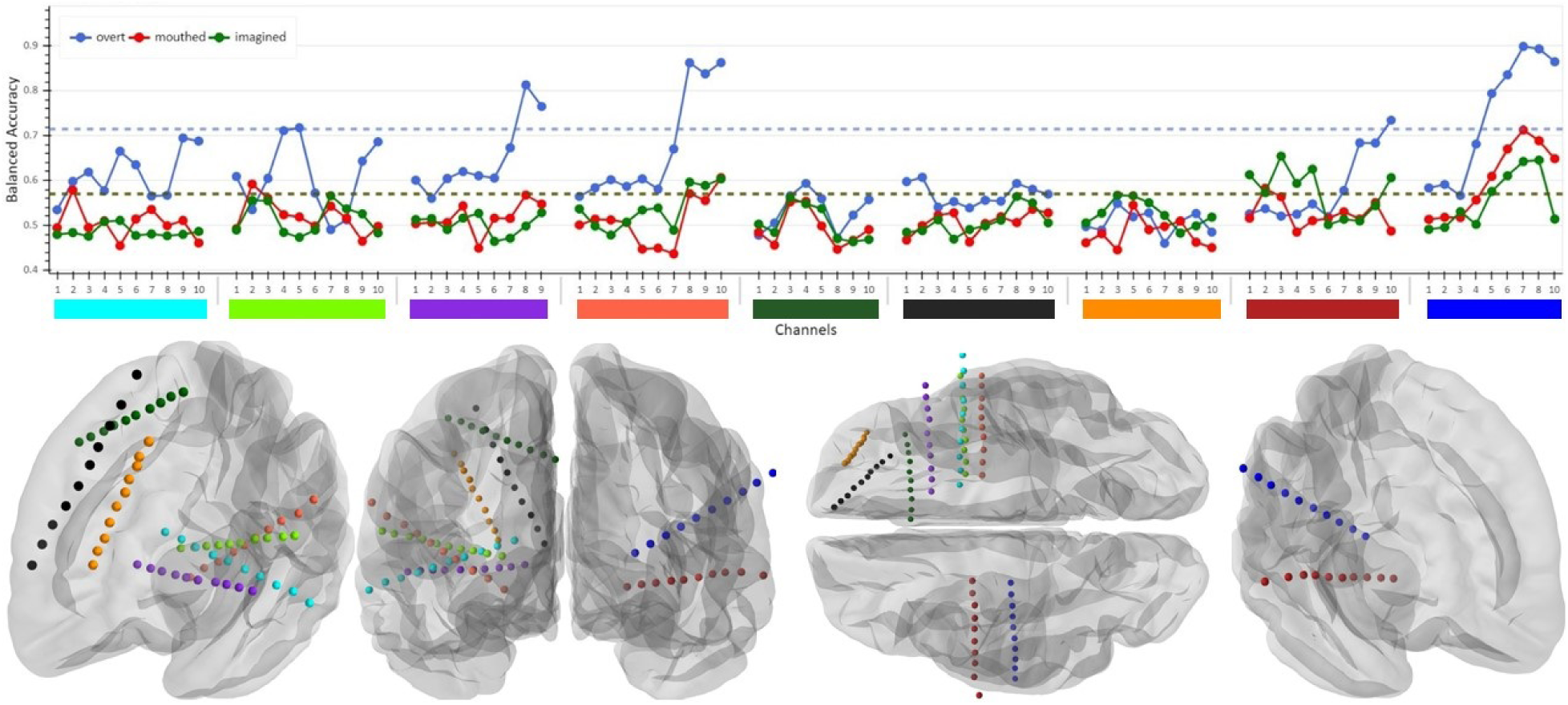
Averaged balanced accuracy of single-channel WM decoding models for all channels for a representative participant (Participant 1). Channels are grouped by shaft, with 1 representing the deepest channel and 10 representing the most superficial channel. The blue, red, and green horizontal dashed lines show the selection thresholds for each mode, averaged across respective folds. Chance-level was approximately 0.5 on average. The color-coded bars below each shaft plot correspond to the colored electrodes in the caudal and ventral views of both hemispheres and frontal views of left and right hemispheres at the bottom of the figure.

Between each mode pair, channels were observed with performance significantly better than chance in both modes (*p <* 0.05, *n* = 10), while having a significantly better performance in the HBO mode than the LBO mode (*p <* 0.05, *n* = 10). For instance, channels in the superior temporal gyrus of both hemispheres (channels 5-8 and 10 on the blue shaft and 7-10 on the light red shaft) show roughly similar above-chance performance for both mouthed and imagined, and significantly better performance for overt (*p <* 0.05, *n* = 10). However, there were also channels that exhibited significantly better in the HBO mode than in the LBO mode (*p <* 0.05, *n* = 10), while showing no significant difference from chance-level performance in the LBO mode (*p >* 0.05, *n* = 10). This is observed for channels in the left middle temporal gyrus (channels 9 and 10 on the cyan shaft) and left inferior temporal gyrus (channels 4, 5, 9, and 10 on the light green shaft), which perform significantly better for overt than mouthed or imagined (*p <* 0.05, *n* = 10), while showing no significant difference from chance-level performance for mouthed and imagined (*p >* 0.05, *n* = 10). Despite a few channels exhibiting slightly higher average performances for imagined than overt and mouthed (i.e., channels 3, 4, and 5 on the red shaft), these are not statistically significant and no channels exhibited significantly better performance in mouthed or imagined than overt.

The averaged balanced accuracy values of the single-channel WM models, normalized for each participant over all three modes, are shown spatially in Fig. 6a. It is observed that most selected channels are clustered in groups of two or more adjacent channels along the same shaft.

**Fig. 6:**
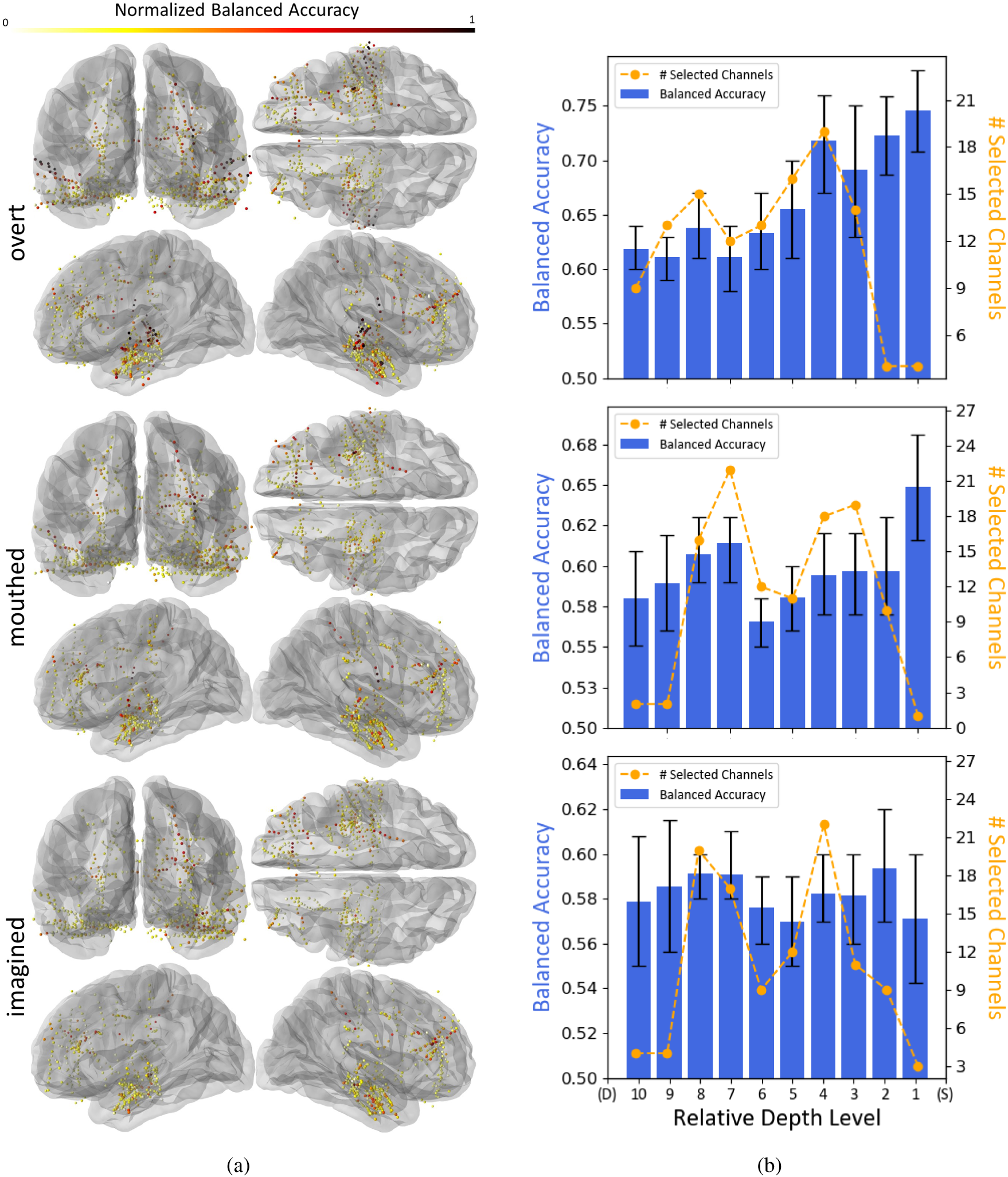
(a) Channels from the single-channel WM analysis having averaged balanced accuracy significantly above chance-level for all participants on an averaged brain model (*p <* 0.05, *n* = 10). The electrodes are colorized based on the averaged balanced accuracy values, which were normalized to 0-1 for each participant over all three modes. (b) Average balanced accuracy across participants and number of selected channels of the single-channel WM decoding models, grouped by relative electrode depth. Selected channels are grouped into ten uniform depth levels based on the center of the Montreal Neurological Institute (MNI) average brain model [43] from the deepest (D) to the most superficial (S) along the respective electrode shafts. The error bars indicate the 95% confidence intervals of balanced accuracy over channels. Chance-level was approximately 0.5 on average.

The overt panel of Fig. 6a shows that relevant channels are predominantly located around the border between the parietal and temporal lobes, including the middle frontal gyrus, superior temporal gyrus and sulcus, and Sylvian parieto-temporal regions. Additionally, some relevant channels are located around the region of the auditory cortex. However, these channels were also selected in one or both of mouthed or imagined, which did not involve auditory feedback. This is consistent with previous studies showing activation in the auditory cortices during imagined speech or imagined hearing, regardless of the presence of an auditory stimulus [36], [44], [45], [24], [46]. Furthermore, these channels were not excluded during the pre-screening for channels exhibiting auditory feedback.

The imagined panel of Fig. 6a also indicates that there are channels, such as those in the left frontal lobe, relevant to imagined that were also selected as relevant to overt and mouthed. However, these channels generally exhibited lower performance compared to the top performing channels of each of these modes. A similar result is observed between overt and mouthed when comparing channels across modes, such as the ones in the right frontal lobe.

In contrast, it can be seen that numerous relevant channels for overt, primarily located in the right and left temporal lobes, were not selected for mouthed and imagined. This is also observed between mouthed and imagined. However, there are several channels located in the right and left temporal lobes, such as channels in the superior temporal gyrus of both hemispheres of Participant 1, that perform well in two or more modes.

To characterize the activity with respect to relative electrode depth, the radial distance between the closest selected channel to the center of the MNI brain model [43] and the furthest selected channel from the center was divided into ten uniformly-spaced levels, forming spherical shells. For each level, the average of the balanced accuracies of the channels located in that level was calculated. Fig. 6b shows the accuracy and number of electrodes corresponding to each relative depth level and mode. The overt results in Fig. 6 show that the best performing channels are comparatively few and near the cortical surface. However, there are a greater number of channels selected at multiple intermediate depths that also exhibit reasonable performance. These observations are also generally consistent for mouthed and imagined.

#### 2) Comparison Across Modes

The results from the single-channel models for the WM and CM paradigms were used to assess and spatially visualize the shared relevance of channels across modes. Channels were organized in three groups according to shared relevance for the nested behavioral output hierarchy of (1) overt, (2) overt-mouthed, and (3) overt-mouthed-imagined. The channels relevant to only overt were selected if the respective performances were significantly above the chance level for the overt WM paradigm and not significantly above chance for either overt-to-mouthed or overt-to-imagined. The channels relevant to both overt and mouthed were selected if they were significantly above chance for both the overt and mouthed WM paradigms, both overt and mouthed CM paradigms, but not significantly above chance for either overt-to-imagined or mouthed-to-imagined. The channels relevant to all three modes were selected if they were significantly above the chance level for all three WM paradigms and all combinations of CM paradigms. Channels were selected based on the respective performance differences in the groups using a threshold of *p <* 0.05.

Fig. 7a shows these three groups of channels from all participants on an average brain model. Channels selected in the overt group show relatively higher performance in the overt panel of Fig. 6. These channels reside in a wide range of cortical and sub-cortical brain regions, including the superior and middle temporal gyrus and superior frontal gyrus, which is in line with previous studies [21], [47], [24], [48], [49], [46], [50].

**Fig. 7:**
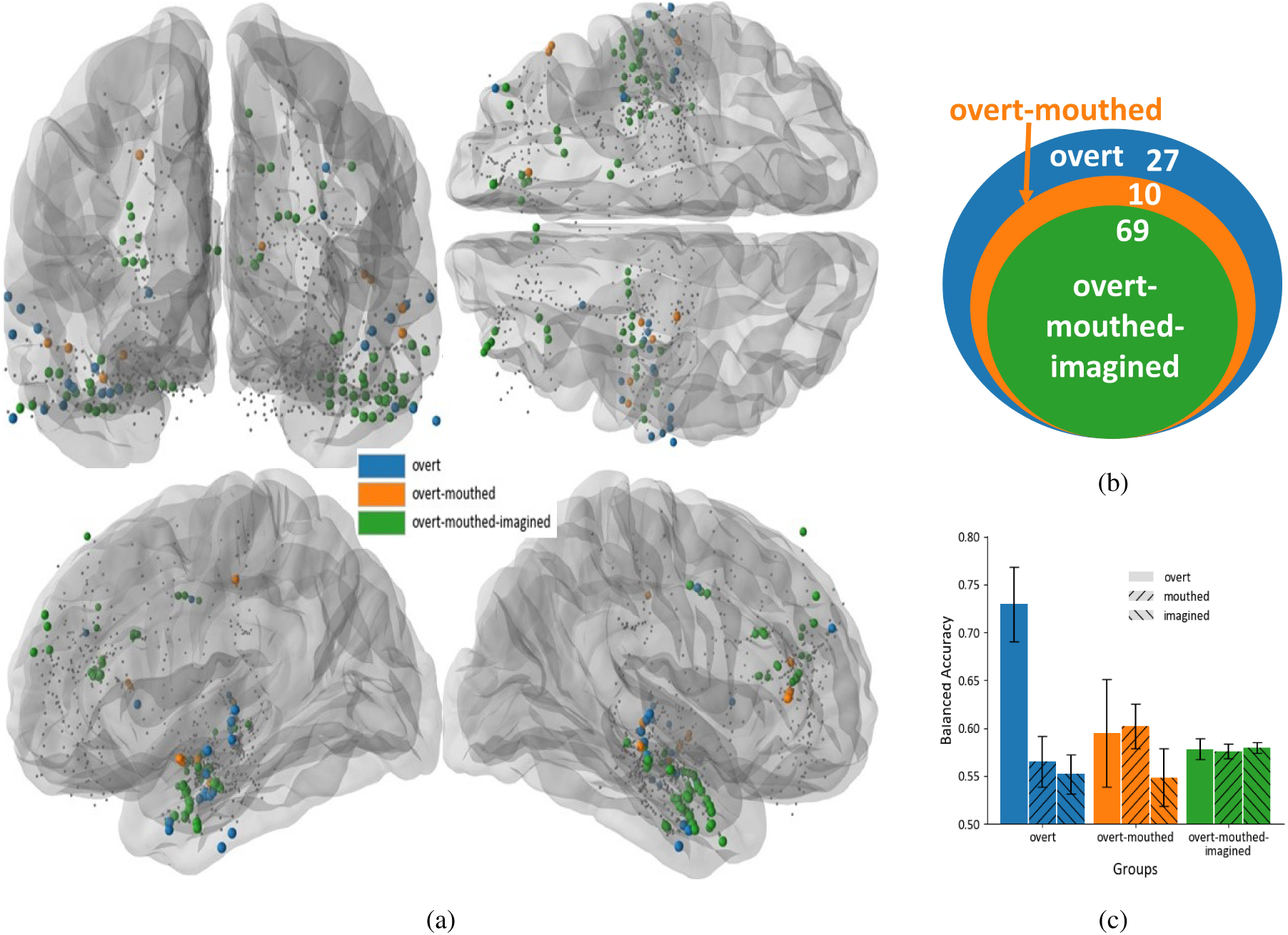
Representations of the hierarchical channel nesting. (a) Channels for all participants on an averaged brain model, color-coded by mode relevance. *Overt-mouthed-imagined* represents the subset of selected channels that are relevant to all three modes. *Overt-mouthed* represents the mutually exclusive subset of channels with processes only relevant to overt and mouthed but not imagined. *Overt* represents the mutually exclusive subset of channels with processes only relevant to overt. Channels not satisfying the relevance criteria are shown as smaller grey points. (b) Venn diagram representing nested hierarchical mode groupings including the number of channels identified in each group, selected from more than 800 channels across all participants. (c) Average balanced accuracy across participants of the single-channel WM decoding models in hierarchical mode groupings. For each grouping, the results are compared across the three modes indicated by the legend using 10-fold non-shuffled cross-validation process for each respective mode. The error bars indicate the 95% confidence intervals. Chance-level was approximately 0.5 on average.

Channels selected in the overt-mouthed group show relatively higher performance in the mouthed panel of Fig. 6. Fewer channels were selected in this group in comparison with the other two groups, which could be due to the narrow coverage of the motor cortices for these participants. These channels are primarily located on or near the motor cortices, right superior frontal gyrus, and right and left inferior temporal gyrus, which is in line with previous studies [51]. Channels selected in the overt-mouthed-imagined group show relatively higher performance in the imagined panel of Fig. 6, which are predominantly on or near the Broca’s area (e.g., left inferior and middle frontal gyrus) and the auditory cortices (e.g., middle and inferior temporal gyrus and sulcus and Sylvian parieto-temporal region), which is in line with previous studies [52], [47], [36], [24], [48], [46], [50]. Selected channels in all three groups reside in both brain hemispheres and in both grey and white matter, which is in line with recent studies that have reported speech-related activity in both hemispheres and both deeper and superficial brain areas during both overt and imagined [2], [52], [53], [54], [46], [50].

It should be noted that channels in the overt group may also contain features shared with the other modes, but these features are relatively less dominant than those unique to overt, as the models trained on overt do not perform significantly above the chance-level on the mouthed or imagined data. This also applies to the overt-mouthed group, which may have features common in all three modes that are likewise relatively less dominant than those unique to overt. Moreover, no channels were found to have significantly better performance in imagined or mouthed than overt. This HBO-LBO hierarchy was also consistent when grouping channels using only mouthed and imagined and comparing their mouthed performances.

Fig. 7b shows a Venn diagram of the nested hierarchical channel groups from Fig. 7a, with relative areas proportional to the number of channels in each group, indicated by the numeric values. Fig. 7c shows the average performance of these channel groups, in a mutually-exclusive fashion (i.e., the colored channels comprise the respective groups, not the nested subsets). For each mutually-exclusive grouping, the results are compared across the three modes indicated by the legend using 10-fold cross-validation process for each respective mode. While channels in all groups performed significantly above chance-level (*p <* 0.0001, *n* = 10), the channels in overt group performed significantly better in the overt mode than channels in overt or the other two groups in any of the three modes (*p <* 0.01, *n* = 10). No significance difference was observed between the performance of the channels in the overt group in mouthed and imagined modes (*p >* 0.05, *n* = 10). No significant difference was observed between the performance of the channels in the overt-mouthed group in overt and mouthed modes (*p >* 0.05, *n* = 10). The performance of the channels in the overt-mouthed group in both overt and mouthed modes was significantly better than the same channels in the imagined mode, the channels in the overt group in mouthed and imagined modes, and the channels in the overt-mouthed-imagined group in overt, mouthed, and imagined modes (*p <* 0.05, *n* = 10). Lastly, no significant difference was observed between the performance of the channels in the overt-mouthed-imagined group in overt, mouthed, and imagined modes (*p >* 0.05, *n* = 10), but each individually was significantly larger than the channels in the overt group in mouthed and imagined modes and the channels of the overt-mouthed group in the imagined mode (*p <* 0.05, *n* = 10).

To directly compare the performance of channels for different modes, performance of each selected channel from Fig. 7a was compared across HBO-LBO mode pairs (i.e., overt-mouthed, overt-imagined, and mouthed-imagined). For each mode-pair and group, all channels selected in at least one mode in the pair based on the respective single-channel WM models were analyzed. Fig. 8 shows a scatter plot of the mode-pair performance of each channel across all participants. Channels are marked according to mode-pair and colorized by nesting grouping. As a visualization aid, bivariate Gaussian distributions were fit to the channels of each nested group. The 90% probability contours of the respective distributions are shown as ellipses in Fig. 8. The average of 10-fold cross-validation performance of the individual channels was generated for each group and the Pearson correlation coefficient was calculated for the performance of each channel in the mode pairs. While a significantly positive correlation (*r* = 0.69 and *p <* 0.0001) was observed between the LBO and HBO axes of overt-mouthed-imagined group, no significant correlation was observed between the LBO and HBO axes of the other two groups (*p >* 0.05).

**Fig. 8:**
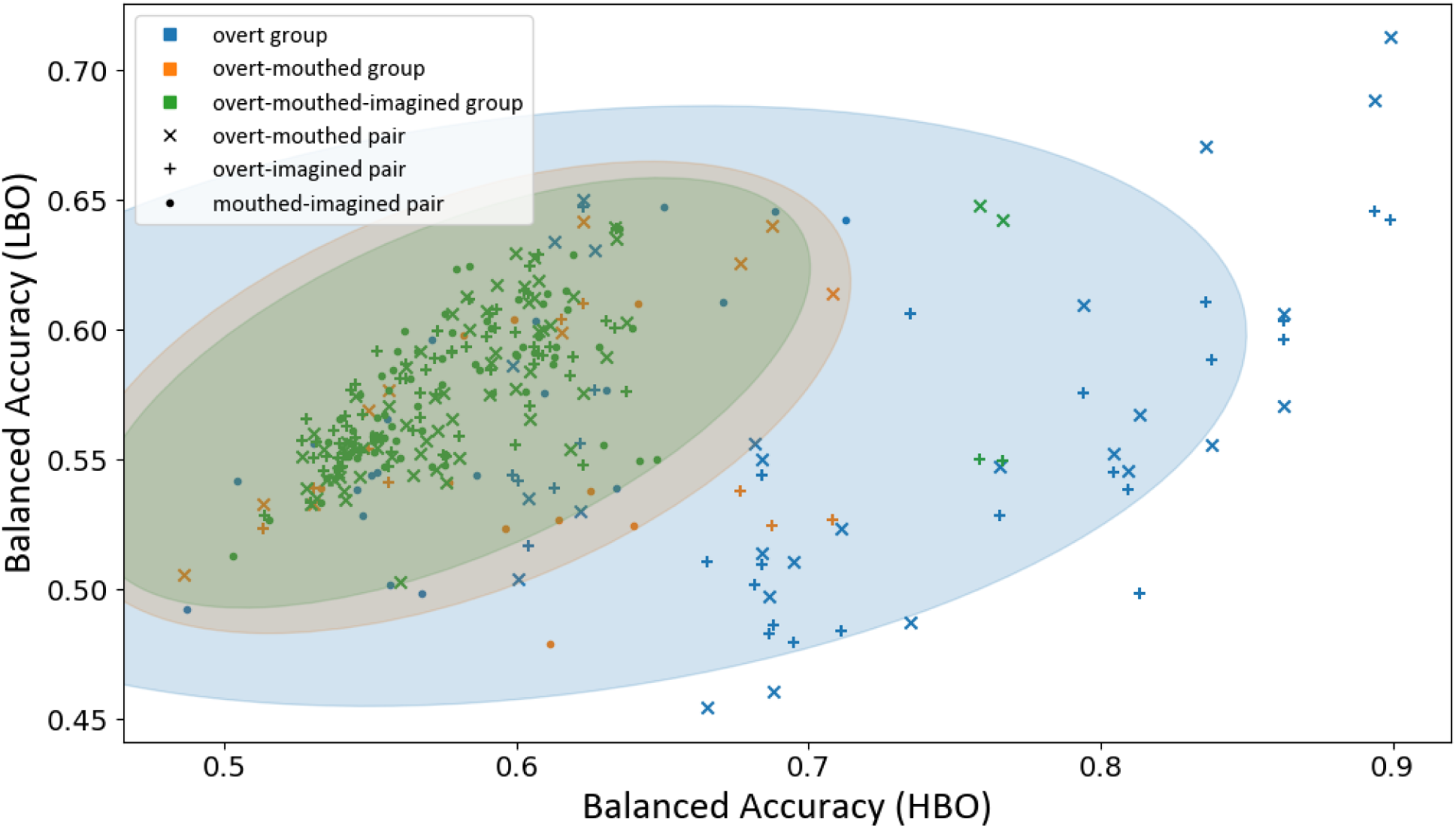
Scatter plot of the mode-pair performance of each selected channel of each group from Fig. 7a across all participants. Channels are marked according to mode-pair and colorized by nesting grouping. As a visualization aid, bivariate Gaussian distributions were fit to the channels of each nested group, and the 90% probability contours are shown as ellipses. Each sample represents the averaged balanced accuracy over the 10-fold cross-validation of two modes of one channel, with the horizontal axis indicating HBO mode and the vertical axis indicating the LBO mode. Chance-level is approximately 0.5 on average.

#### 3) Spectro-temporal Characterization

Fig. 9 illustrates the absolute value of normalized feature weights of the single-channel WM models for each mode, averaged over the selected channels of the channel groups from Fig. 7a. While these weights do not directly represent the spectro-temporal brain patterns associated with the decoding models, they do convey the relative contributions of spectro-temporal features to the models. As expected from previous studies, features temporally closer to the frame being decoded have a greater contribution to the models [2]. A strikingly similar pattern is observed between the weights of the overt-mouthed-imagined group across the three modes. Such similarities are also observed for the overt-mouthed group in the overt and mouthed modes. While, as expected, broadband gamma was a prominent feature, it was observed that the lower frequency bands also provide important contributions to the models. This also supports previous studies that have shown alpha band to be promising for distinguishing movement from rest [55] and *speech* from *non-speech* [2].

**Fig. 9:**
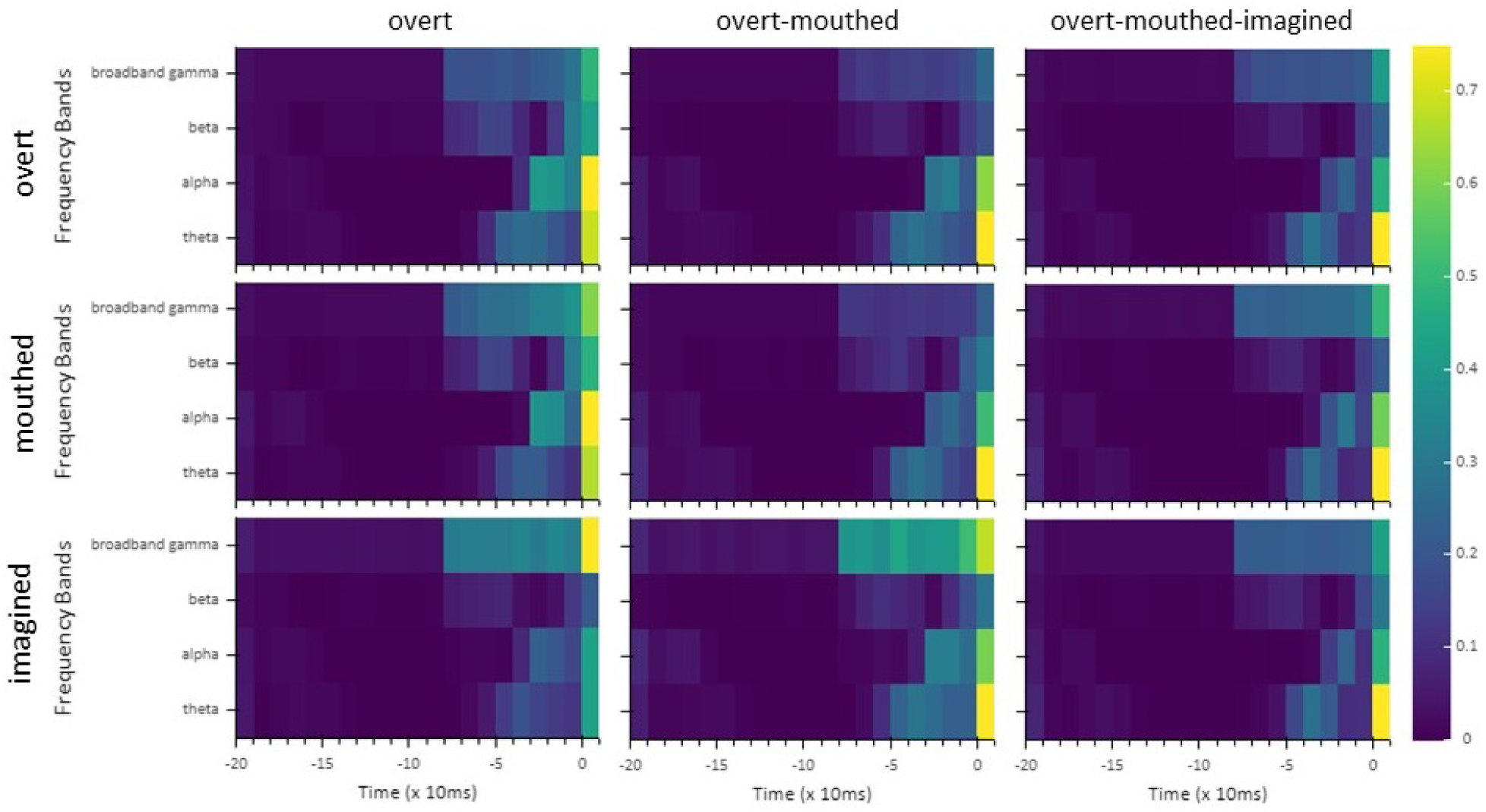
Average of absolute value of normalized decoding model weights across 10-folds of the channel groups from Fig. 7a. Zero on the horizontal axis indicates the start of the audio frame.

### B. Models: Within-Mode and Cross-Mode

Fig. 10 shows the averaged balanced accuracy in the multi-channel WM and CM models across participants. All multi-channel WM and CM models performed significantly better than chance-level (*p <* 0.05, *n* = 10 and 50 for WM and CM models, respectively), except for the overt-to-mouthed and overt-to-imagined models of Participant 3. Overt-to-overt models performed significantly better than all other models (*p <* 0.0001, *n*_1_ = 10 and *n*_2_ = 10 or 50), which may be attributed to the more precise labeling of the overt trials. Models trained on mouthed performed significantly better than all other models (*p <* 0.01, *n*_1_ = 10 or 50 and *n*_2_ = 10 or 50) on both mouthed (mouthed-to-mouthed models) and on overt (mouthed-to-overt models), except imagined-to-imagined models. However, no significant difference was observed between the performance of mouthed-to-mouthed, imagined-to-imagined, and mouthed-to-overt models (*p >* 0.05, *n* = 10, 10, and 50, respectively). Overt-to-mouthed models performed significantly better than overt-to-imagined models (*p <* 0.0001, *n* = 50), and mouthed-to-overt models performed significantly better than imagined-to-overt models (*p <* 0.01, *n* = 50).

**Fig. 10:**
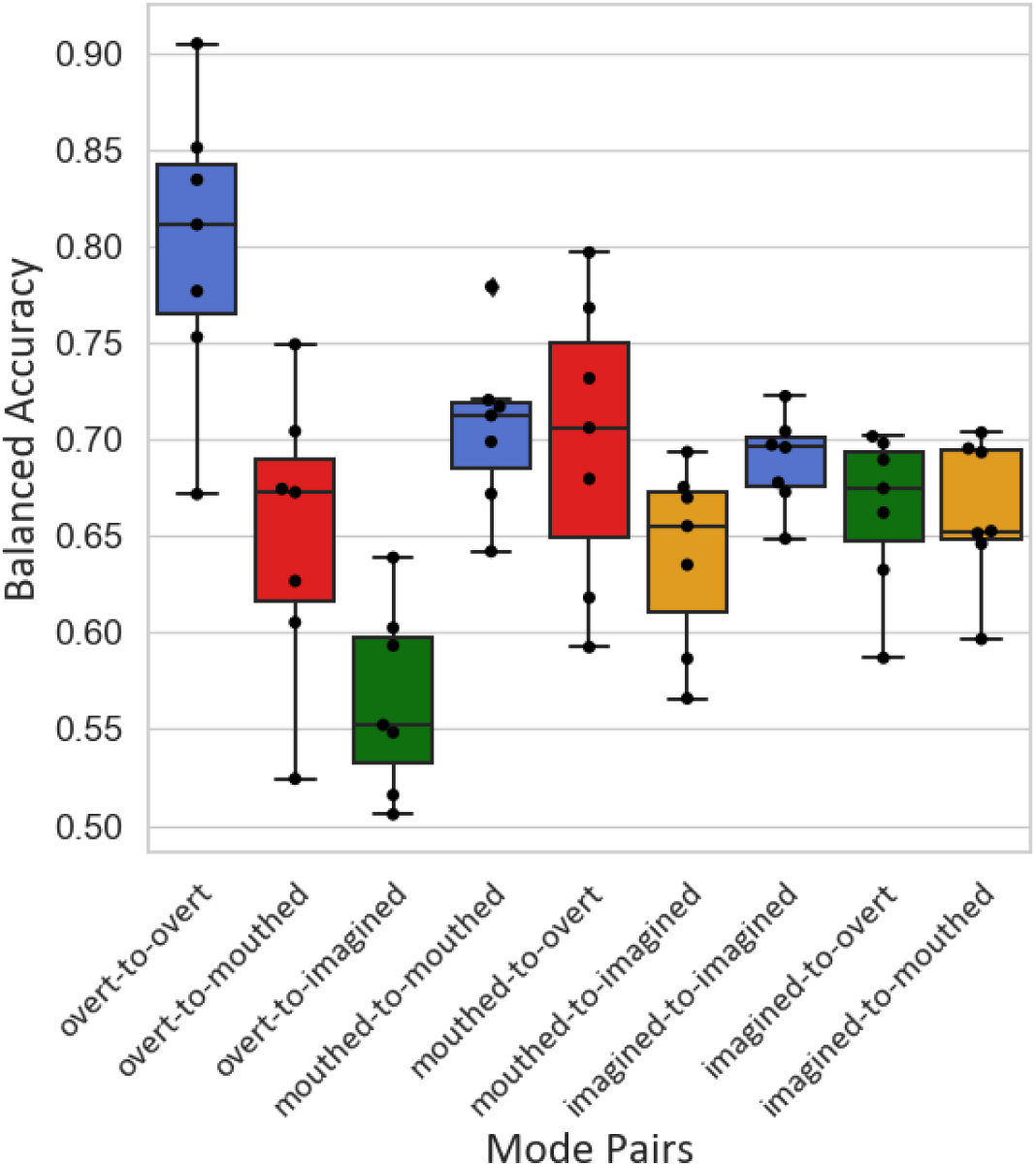
Box plot of the average balanced accuracy of the multi-channel WM and CM models across participants. Blue boxes represent the WM models. Red boxes represent both train-test combinations of the CM models for overt and mouthed. Green and orange boxes represent the CM models for the combinations of overt/imagined and mouthed/imagined, respectively. The horizontal line within each box shows the median, while the extents of the boxes represent the first (Q1) and third (Q3) quartiles. The whiskers extend from the box to 1.5 times the inter-quartile range (IQR). Each dot represent a data point from an individual participant that lies between the two 1.5IQRs and the outliers are indicated with diamonds. Chance-level was approximately 0.5 on average.

In contrast, the performance of the mouthed-to-imagined models was only significantly better than the overt- to-imagined models (*p <* 0.001, *n* = 50), and the performances of both of these two models were significantly worse than all other multi-channel WM and CM models (*p <* 0.05, *n*_1_ = 10 and *n*_2_ = 10 or 50). Moreover, the performances of the imagined-to-overt and imagined-to-mouthed models were significantly better than the performances of the overt-to-imagined and mouthed-to-imagined models, respectively (*p <* 0.05, *n* = 50). However, no significant difference was observed between the performances of the imagined-to-overt and imagined-to-mouthed models (*p >* 0.05, *n* = 50).

The performance of the imagined-to-overt models was significantly worse than both the overt-to-overt and mouthed-to-overt models (*p <* 0.0001, *n* = 50, 10, and 50, respectively). Also, the performance of the imagined-to-mouthed models was significantly worse than the mouthed-to-mouthed models (*p <* 0.01, *n*_1_ = 50 and *n*_2_ = 10), but no significant difference was found in comparison to the overt-to-mouthed models (*p >* 0.05, *n* = 50). Mouthed-to-overt and imagined-to-overt models performed significantly better than the overt-to-mouthed and overt-to-imagined models, respectively (*p <* 0.05, *n* = 50). Imagined-to-mouthed models performed significantly better than the mouthed-to-imagined models (*p <* 0.05, *n* = 50).

It should be noted that, for each participant and mode, the selected channels from Section II-H.2 exhibited largest contributions to the multi-channel models.

### C. ch Activity Detection Performance

Fig. 11a shows the distributions of proportions of speech activity detection in each of multi-channel WM and CM models in comparison with the actual proportion of speech based on the true or approximated labels over all participants. The mean of the detected speech proportions for all multi-channel WM models was significantly larger than the mean of the speech proportions based on the actual labels (*p <* 0.0001, *n* = 50). For the WM models, the proportions of detected speech for overt-to-overt was significantly smaller than both mouthed-to-mouthed and imagined-to-imagined (*p <* 0.05, *n* = 50); however, no significant difference was found between mouthed-to-mouthed and imagined-to-imagined (*p >* 0.05, *n* = 50).

**Fig. 11:**
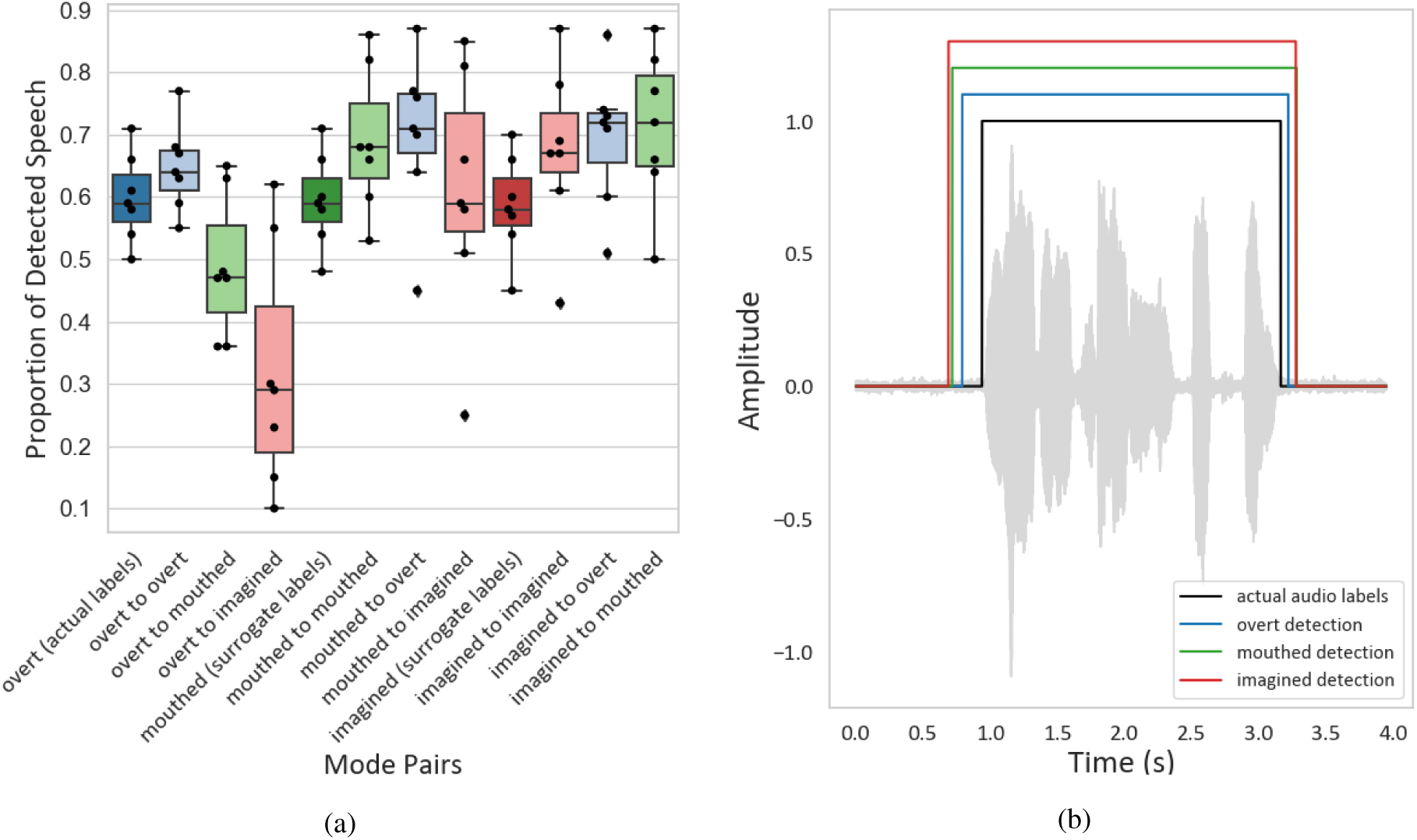
(a) Average proportions of decoded speech vs. actual/surrogate speech labels for all participants and modes for all multi-channel WM and CM models. The bold-shaded boxes represent the speech proportions based on the actual/surrogate labels. For reference, the colors of lighter-shaded boxes are coordinated with colors of the actual/surrogate labels of the respective test modes. Refer to Fig. 10 for a description of the box plot properties. (b) A representative speech activity detection model outputs vs. the actual audio labels for one sentence from a representative participant (Participant 4).

For all WM models, the detected speech windows ended significantly later than when the actual or estimated speech ended (*p <* 0.0001, *n* = 50), while for all mouthed-to-mouthed and imagined-to-imagined WM models, the detected speech windows started significantly earlier than when the actual or estimated speech started (*p <* 0.01, *n* = 50). For all overt-to-overt WM models, the detected speech windows started significantly later than the detected speech windows of mouthed-to-mouthed and imagined-to-imagined WM models (*p <* 0.01, *n* = 50), while no significant difference was observed between these window starts and when the actual speech started (*p >* 0.05, *n* = 50). Fig. 11b shows the speech detection versus actual speech for overt, mouthed, and imagined of a representative participant (Participant 4).

For all multi-channel CM models, except the overt-to-mouthed and overt-to-imagined models, the mean of the detected speech proportions was significantly larger than the mean of the speech proportions based on the actual or estimated labels (*p <* 0.0001, *n* = 50). For the overt-to-imagined models, the mean of the detected speech proportions was significantly lower than the mean of the speech proportions based on the estimated labels (*p <* 0.01, *n* = 50). This can be related to the relatively lower performance of these two models as depicted in Fig. 10. No significant difference was observed between the mean of the detected speech proportions of overt-to-mouthed models and the mean of the speech proportions based on the estimated labels (*p >* 0.05, *n* = 50).

## IV. Discussion

This study used sEEG data collected during overt, mouthed, and imagined speaking conditions to identify common neural features and relationships across conditions using a speech activity detection paradigm. The relevant features were found to occur near speech-onset, across all frequency bands examined as shown in Fig. 9, which is in line with previous studies [2].

### A. Nested Behavioral Hierarchy: Single Channel Models

Recent studies in Neurolinguistics have offered evidence for the existence of a nested hierarchy in the brain activity associated with different speech modes, formed from highest behavioral output to lowest behavioral output [13], [14], [56], [57]. Facial micromovements during imagined speech, commonly assumed to be a byproduct of short-circuited motor signals and induced activity in language-associated brain areas (e.g., Broca’s and Wernicke’s areas) during both overt and imagined speech are among the evidence supporting this hypothesis [13], [46], [58], [59], [60]. It is hypothesized that imagined speech is an abbreviation of overt speech, suggesting that brain processes relevant to imagined speech are also involved during overt speech, whereas overt speech involves additional brain processes beyond imagined speech - likely associated with articulatory planning, articulation, sound production, and possibly aspects of perceptual feedback.

This study provides evidence that the channels relevant to different speech modes generally form nested hierarchical subsets from highest behavioral output to lowest behavioral output. Specifically, channels relevant to imagined were found to be a subset of those relevant in mouthed, while those relevant for mouthed were a subset of those relevant for overt. The subset of relevant channels for mouthed and overt that is mutually exclusive with imagined likely represents activity related to direct control of the speech articulators modulated in both modes. The channels exclusive to overt are presumed to be related to brain activities present exclusively for overt, including perceptual feedback, articulatory planning, articulatory motor executions, and/or sound production. The perceptual feedback likely represents indirect perceptual activity since activity from direct auditory feedback was excluded during screening. The subset of channels for all modes is hypothesized to represent the common core of activity for general speech activity production.

When examining individual channels across modes, as shown in Fig. 7, the nested nature of the channels within the mode hierarchy is apparent. The majority of relevant channels were shared amongst the three modes, while only about ten percent were unique to overt and mouthed but not imagined. The channel subset relevant to imagined speech was found to reside in bilateral frontal and temporal regions, which is consistent with prior ECoG work indicating that overt and imagined produce neural activity in both right and left cortical hemispheres [47]. Furthermore, these activations occurred at various bilateral depths as indicated in Fig. 6b. This is consistent with previous studies showing channels from both grey and white matter contributing to decoding models [8], [2], [50], [55], further demonstrating the relevance of deeper structures and white matter for speech decoding.

Roughly a third of the relevant channels were unique to overt, which predominantly resided in more superficial temporal regions, despite pre-screening channels for auditory feedback. Notwithstanding the absence of auditory feedback, activations on or around the auditory cortex were observed for mouthed and imagined. This is consistent with prior studies using overt and imagined speech and has been hypothesized to be related to inner speech rehearsal [14], [15], [47], [61], [36], [60], [62].

Fig. 7c and Fig. 9 further support the nested behavioral hierarchy by comparing the model performances and respective feature weights of the nested channel groups across modes. The overt-mouthed-imagined group exhibits highly consistent performance across the three modes, while performance is degraded for the other groupings when evaluated on the LBO modes. While the relevant spectro-temporal features of the overt-mouthed-imagined group are quite similar across all three modes, the features of the overt-mouthed group for the overt and mouthed are also similar and noticeably different from imagined. Furthermore, differences between the feature of the overt group are observed between overt and the two LBO modes.

This hierarchy, with respect to relative decoding performance of relevant channels between mode pairs, is also observed from Fig. 8. Nearly all channels yield an above chance performance for each mode in the pairs, except for a select group of channels in the lower portion of the plot that perform well for overt but not the other modes. This indicates that nearly all channels with above-chance performance in the LBO mode also performed above chance in the HBO mode, while the inverse does not hold. The majority of channels in the overt-mouthed-imagined group and the mouthed-imagined pair of the overt group are clustered within the same range on the LBO and HBO axes (i.e., 0.5-0.65), suggesting that these channels roughly yield comparable performance for both modes. However, the majority of the overt-mouthed and overt-imagined pairs of the overt group reside toward the right of the plot, with the HBO axis (overt mode) having noticeably larger values, showing that these channels have a more dominant and potentially unique activity during overt compared to the other two modes. This is relationship also apparent in the overt group of Fig. 7c. A weaker but similar trend is observed among the overt-mouthed group, with majority of values in the same range on the LBO and HBO axes (i.e., 0.5-0.65) and some located toward the bottom right of the plot, with the HBO axis (overt or mouthed) having noticeably larger values than the LBO axis (imagined). This is also apparent in overt-mouthed group of Fig. 7c.

### B. Further Evidence: Multi-Channel Models

These findings are relevant for understanding why decoding models successfully trained and tested using overt speech tend to be poor at generalizing to imagined speech, as shown in Fig. 10. This figure also shows that there is a consistent decrease in performance when training on an HBO mode and testing on an LBO mode compared to the inverse, and this decrease is more pronounced for larger differences in the behavioral output hierarchy. It is also observed that the imagined models perform consistently well across modes, while the overt-to-imagined performs poorest amongst the combinations. This further suggests that the relevant channels for the imagined models also represent speech processes present for mouthed and overt, whereas the other relevant channels from overt (likely associated with articulation and possibly aspects of perceptual feedback) do not extend to imagined. When interpreting these results, it is important to note that although the relevant LBO channels are available when training the HBO models, they do not appear to be selected or weighted in a way to generalize to the LBO modes.

While the present results offer strong evidence for the nested behavioral output hierarchy, some other studies suggest this may be an oversimplification and imagined speech can be more than just an abbreviation of overt speech processes [13], [14], [56], [57], [52], [63]. These prior studies indicate that imagined speech may involve different linguistic processes than those relative to overt speech, or may contain unique brain processes (e.g., inhibitory activity) that are not involved in overt speech such as more prominent activity in the middle frontal gyrus, left and right temporal gyrus, left supramarginal gyrus, left superior frontal gyrus, and in various regions of white matter [13], [14], [52], [64], [40]. Nonetheless, other studies have proposed that imagined speech may be an abbreviation of overt speech processes, but further investigation is required to verify the precise mechanisms at the linguistic and motor levels [13], [57]. The present study is limited by the nature and availability of sEEG recordings from a relatively low number of participants with sparse and inconsistent electrode coverage. While this coverage is not designed or ideal for speech decoding, it does provide important insights regarding previously unexplored features for this purpose.

### C. Speech Activity Detection Performance

While the use of the speech activity detection model provides a very coarse labeling for the actual and surrogate speech, it has yielded statistically above-chance performing models across modes, thus providing a solid and simplified basis for exploring and comparing the models and relevant features. Fig. 11a shows that the proportions of detected speech tend to be overestimated when training using the surrogate labels. This is likely due to the inherent variability when applying surrogate labels. In contrast, the proportion of detected speech is underestimated when training on the actual (overt) labels and testing on the surrogate labels, and this effect is more pronounced for decreasing behavioral output. The detected proportions are strikingly consistent when training on imagined and testing across modes. This further suggests the existence of a nested behavioral output hierarchy.

It was observed that the detected windows generally lead the actual speech-onset and lag the speech-offset (Fig. 11b), resulting in a higher false positive rate than false negative rate. This is again likely due to the inherent variability of the surrogate labels, but nevertheless may be desirable in practical application where the primary goal is to reliably detect the intention to speak.

## V. Conclusion

The main objective of this study was to elucidate neural features associated with imagined speech to inform the development of imagined-speech neuroprostheses. This was achieved by comparing neural features and associated speech activity detection decoding model performance across three speech modes with varying degrees of behavioral output. The results suggest that the relevant channels can be organized in a nested hierarchy according to the degree of behavioral output, with the overt mode encompassing all relevant channels across modes, the relevant channels from the mouthed mode being a subset of overt, and the relevant channels from the imagined mode being a subset of mouthed. This nested hierarchy suggests that there may be a common neural substrate of related speech processes that progressively extends with increasing behavioral output. These findings also provide important insights toward the design and development of imagined speech decoding models based on available overt speech data. Additionally, through the acquisition of sEEG, relevant channels across modes were found beyond the cortex, bilaterally at various depths, in both grey and white matter. This provides further evidence that deeper structures are relevant and may be beneficial in the development of improved speech decoding models. These findings also show that, with proper consideration and treatment, recordings of overt speech can serve as viable surrogates for generating imagined-speech decoding models. Given the limitations of sEEG recordings in terms of coverage and patient accessibility, additional work is needed to further characterize and understand the neural activity relationships across speaking modes. While the speech activity detection model provides a simplified framework for comparison, it is envisioned that these findings can be extended to more sophisticated imagined speech decoding schemes to reveal more nuance to the features and relationships.

## Data and code availability

The data for the present study is available through reasonable request to the corresponding author. The code from this study will be made publicly available via a suitable open repository when the peer-reviewed manuscript is published.

## Author contributions

P.S. and D.K. designed the study, and C.H., T.S., and D.K. designed the experiments. S.R. and J.S. were involved in data collection, and P.S. analyzed the data. P.S. and D.K. discussed the results and wrote the manuscript, and all authors contributed to review and editing of the manuscript. D.K., T.S., and J.S. acquired financial support for the project.

